# Conserved and ubiquitous expression of piRNAs and PIWI genes in mollusks antedates the origin of somatic PIWI/piRNA expression to the root of bilaterians

**DOI:** 10.1101/250761

**Authors:** Julia Jehn, Daniel Gebert, Frank Pipilescu, Sarah Stern, Julian Simon Thilo Kiefer, Charlotte Hewel, David Rosenkranz

**Author notes:** contributed equally (Julia Jehn, Daniel Gebert, Frank Pipilescu). Correspondence to: David Rosenkranz.

## Abstract

PIWI proteins and a specific class of small non-coding RNAs, termed Piwi interacting RNAs (piRNAs), suppress transposon activity in animals on the transcriptional and post-transcriptional level, thus protecting genomes from detrimental insertion mutagenesis. While in vertebrates the PIWI/piRNA system appears to be restricted to the germline, somatic expression of piRNAs directed against transposons is widespread in arthropods, likely representing the ancestral state for this phylum. Here, we show that somatic expression of PIWI genes and piRNAs directed against transposons is conserved in mollusks, suggesting that somatic PIWI/piRNA expression was already realized in an early bilaterian ancestor. We further describe lineage specific adaptations regarding transposon composition of piRNA clusters and show that different piRNA clusters are dynamically expressed during oyster development. Finally, bioinformatics analyses suggest that different populations of piRNAs participate in the ping-pong amplification loop in a tissue specific manner.

## Introduction

In virtually all animals, PIWI proteins protect germ cells from the steady threat of mobile genetic elements, so-called transposons (Thomson and Lin 2009, Iwasaki et al. 2015). Based on sequence complementarity to their target transcripts, 23-31 nt non-coding RNAs, termed PIWI-interacting (pi-) RNAs, function as guide molecules for PIWI proteins that endonucleolytically slice snatched targets. Besides post-transcriptional transposon silencing, PIWI proteins and piRNAs can trigger the establishment of repressive epigenetic DNA or chromatin modifications, thus inducing efficient transposon silencing on the transcriptional level (Reuter et al. 2011, Giacomo et al. 2013, Pezic et al. 2014, Manakov et al. 2015).

Analysis on piRNA pathways in representatives of many animal taxa have unveiled a great diversity of lineage specific adaptations, challenging the universal validity of insights obtained from model organisms (Grimson et al. 2008, Houwing et al. 2008, Das et al. 2008, Li et al. 2013, Lim et al. 2014, Ha et al 2014, Hirano et al. 2014, Gebert et al. 2015, Roovers et al. 2015, Rosenkranz et al. 2015, Madison-Villar et al. 2016, Praher et al. 2017, Lewis et al. 2018). For a long time, PIWI proteins and piRNAs were thought to be dispensable for female germ cell development in mammals until it became clear that the model organisms mouse and rat represent an exception from the mammalian rule (Flemr et al. 2013, Roovers et al. 2015). Similarly, evidence for a gene regulatory role of piRNAs (Zhang et al. 2015, Gebert et al. 2015, Russel et al. 2017) and their widespread somatic expression in many animals (Palakodeti et al. 2008, Perrat et al. 2013, Nandi et al. 2016, Jones et al. 2016, Lewis et al. 2018) have eroded the dogma that the piRNA pathway is restricted to the germline, being exclusively responsible for silencing of transposons.

Beyond transposon control, piRNAs are essential for regeneration and stem cell maintenance in the flatworm *Schmidtea mediterranea* (Palakodeti et al. 2008), they provide adaptive immunity against virus infections in *Aedes aegypti* (Miesen et al. 2015), are responsible for sex determination in *Bombyx mori* (Kiuchi et al. 2014), memory-related synaptic plasticity in *Aplysia californica* (Rajasethupathy et al. 2012) and cleave mRNA in pig, mouse and human (Zhang et al. 2015, Gebert at el 2015).

Despite the more than eighty thousand living molluskan species (Rosenberg 2014) there exists only one description of PIWI proteins and piRNAs for this taxon, based on experiments in the sea slug *Aplysia californica* (Rajasethupathy et al. 2012), making any conclusions on conserved or lineage-specific features of the PIWI/piRNA system in mollusks impossible. In order to overcome this lack of information, we have reconstructed the evolution of PIWI genes in mollusks based on 11 sequenced genomes. We performed quantitative real-time PCR experiments to analyze the expression patterns of the identified PIWI paralogs across a representative set of tissues from the great pond snail *Lymnaea stagnalis* (*L. stagnalis*) and the pacific oyster *Crassostrea gigas* (*C. gigas*). We applied high-throughput sequencing of small RNAs from *L. stagnalis* to verify the presence of piRNAs in germline and muscle tissue. We further reanalyzed published small RNA sequence data from *C. gigas* to characterize the dynamic expression of piRNAs from distinct piRNA clusters during oyster development. Finally, we used bioinformatics approaches to show that different piRNA populations participate in the ping-pong amplification loop in a tissue specific manner.

## Results

### The molluskan PIWI gene repertoire

A number of previously published gene tree reconstructions of PIWI family members suggest that *Drosophila* Ago3 and deuterostomian Piwi-like genes derived from an ancestral gene that was present in the common ancestor of today-living bilaterian species (Seto et al. 2007). Since the number of PIWI paralogs differs across model organisms, we first wanted to characterize the PIWI protein equipment of sequenced mollusks to infer the ancestral state and subsequent evolution of PIWI paralogs in the molluskan clade. We used available PIWI protein sequence data from six molluskan species (*Biomphalaria glabrata*, *Aplysia californica*, *Crassostrea gigas*, *Crassostreas virginica*, *Mizuhopecten* yessoensis, *Octopus bimaculoides*) and further manually annotated PIWI genes based on five publicly available but yet unannotated genomes (*Lymnaea stagnalis*, *Radix auricularia*, *Lottia gigantea*, *Bathymodiolus platifrons*, *Pinctada martensii*). We found that the PIWI family members Piwil1 and Piwil2 are conserved in mollusks and orthologous to Piwil1 and Piwil2 in vertebrates, suggesting a duplication event in an early bilaterian ancestor with subsequent loss of Piwil1 in arthropods (Figure 1A). While we did not observe further gene duplication events within the molluskan Piwil2 clade, several duplication events are present in the Piwil1 clade resulting in two Piwil1 paralogs in *Bathymodiolus platifrons* and even three Piwil1 paralogs in *Lymnaea stagnalis* and *Radix auricularia*. Generally, PIWI gene duplication events are in line with the previously described erratic evolution of PIWI family genes in arthropods (Lewis et al. 2016, Dowling et al. 2016, Lewis et al. 2018). Noteworthy, it was also a successive duplication of Piwil1 on the eutherian lineage that gave rise to Piwil3 (with subsequent loss on the murine lineage) and Piwil4 (Sasaki et al. 2003, Murchison et al. 2008, Figure 1A).

### Ubiquitous expression of PIWI genes in *Lymnaea stagnalis* and *Crassostrea gigas*

To investigate the expression of PIWI genes in mollusks we chose two representative species, the pacific oyster *Crassostrea gigas* (*C. gigas*, Bivalvia) showing no Piwil1 duplication, and the great pond snail *Lymnaea stagnalis* (*L. stagnalis*, Gastropoda), featuring three predicted Piwil1 paralogs (Figure 1A). We performed quantitative real-time PCR for each PIWI paralog on a representative set of tissues from both species.

For the hermaphroditic great pond snail *L. stagnalis* we measured PIWI expression on the mRNA level in the reproductive tract, comprising both male and female gametes, foot muscle, lung and brain. Significant expression was detectable for Piwil1 and particularly Piwil2, while the Piwil1 duplicates Piwil1b and Piwil1c are expressed only at very low levels (Figure 1B and 1C). As expected, we observed the highest expression of Piwil1 and Piwil2 in the reproductive tract. However, both genes are significantly expressed in the other analyzed tissues as well, reaching 15%, 62% and 21% of germline expression for Piwil1 in brain, muscle and lung, respectively, and 12%, 36% and 53% of germline expression for Piwil2 in brain, muscle and lung, respectively (Figure 1D).

Next, we turned to the dioecious pacific oyster *C. gigas*, were we measured PIWI expression on the mRNA level in the male gonad, labial palps, gill, adductor muscle and mantle. We detected significant expression of Piwil1 and Piwil2 across all analyzed tissues, particularly in gonadal tissue (Figure 1E and 1F). In relation to gonadal expression, Piwil1 and Piwil2 are expressed in levels ranging from 21% (Piwil1 in labial palps) to 111% (Piwil2 in adductor muscle, Figure 1G). The observed expression patterns suggest that a functional PIWI machinery acting in the soma and the germline is conserved in mollusks. Considering the somatic expression of PIWI proteins and piRNAs in many arthropod species (Lewis et al. 2018), it is parsimonious to assume that somatic PIWI/piRNA expression represents the ancestral state that was established in an early bilaterian ancestor.

**Figure 1.**
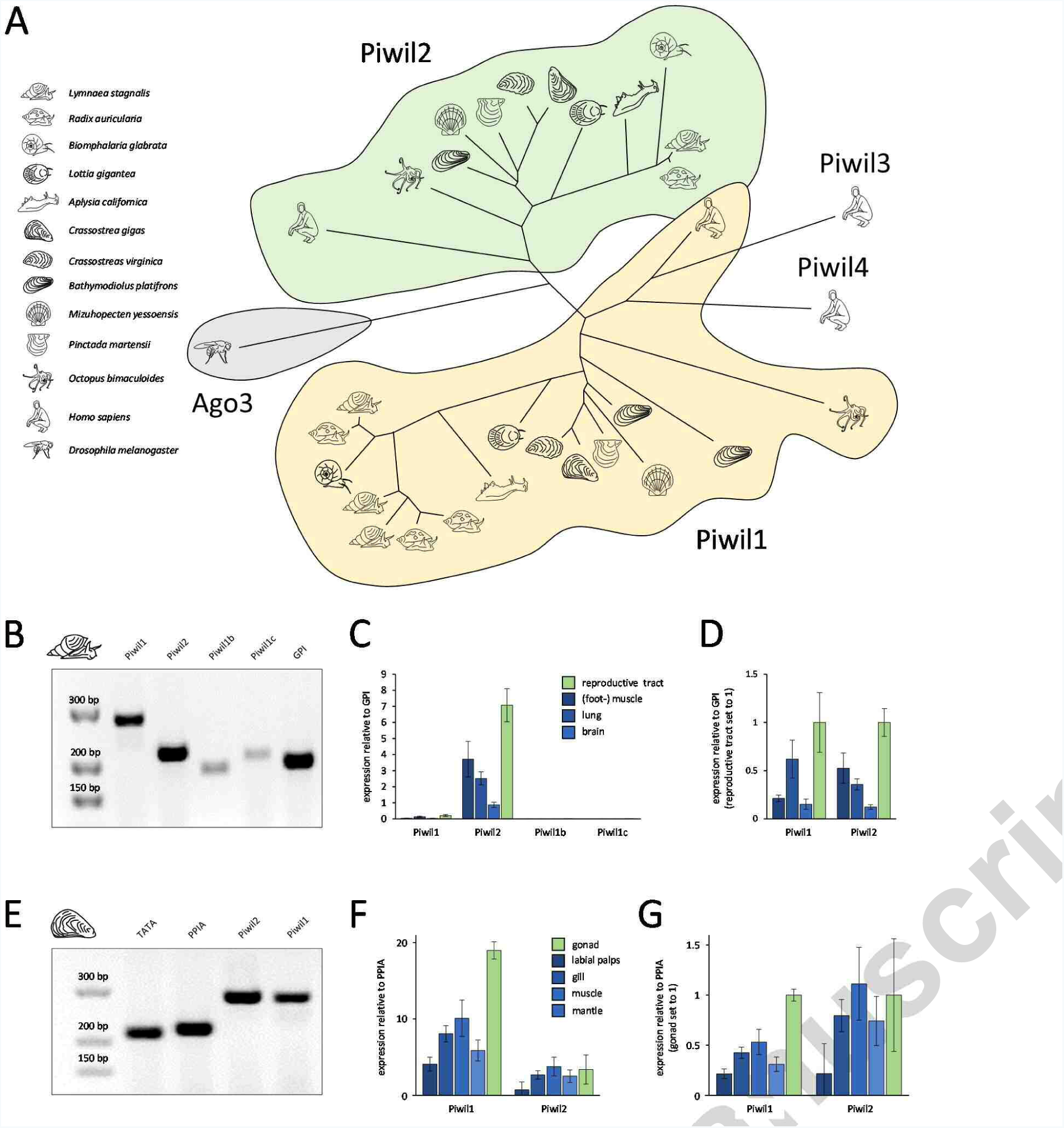
Evolution and expression of PIWI genes in mollusks. (A) PIWI gene tree reconstruction of molluskan PIWI genes. (B) Control PCR with PIWI paralog specific primers and *L. stagnalis* cDNA from the reproductive tract. (C) RT-qPCR results for PIWI paralog expression in different tissues of *L. stagnalis*, normalized by the expression of the housekeeping gene GPI. (D) PIWI paralog expression in different tissues of *L. stagnalis*, normalized by the expression of the housekeeping gene GPI, values from reproductive tract set to 1. (E) Control PCR with PIWI paralog specific primers and *C. gigas* cDNA from the adductor muscle. (F) RT-qPCR results for PIWI paralog expression in different tissues of *C. gigas*, normalized by the expression of the housekeeping gene PPIA. (G) PIWI paralog expression in different tissues of *C. gigas*, normalized by the expression of the housekeeping gene PPIA, values from male gonad set to 1.

### piRNAs in *Lymnaea stagnalis* muscle and reproductive tract

In order to characterize molluskan piRNAs, we sequenced small RNA transcriptomes from *L. stagnalis* extracted from the reproductive tract and (foot-) muscle, since muscle tissue was found to exhibit the highest somatic PIWI expression in both *L. stagnalis* and *C. gigas*. The sequence read length profiles of both probes show a maximum for 21 nt RNAs, with a considerable amount of 22 nt RNAs being present in the muscle, but not in the However, the share of 3’ tRFs in the reproductive tract reproductive tract. We further observed a smaller is nearly twice as high compared to muscle (18% and fraction of RNAs in the range of 24-29 nt in both 10%, respectively, Figure 2B). Recently, 3’ tRFs were probes (Figure 2A). sRNA sequence annotation with found to silence Long Terminal Repeat (LTR) unitas (Gebert et al. 2017) revealed a similar retrotransposons in mouse stem cells by targeting proportion of different sRNA classes in the two their functionally essential and highly conserved probes, with miRNAs accounting for 46% and 51% of primer binding sites (Schorn et al. 2017). The reads in the reproductive tract and muscle, remarkable amount of 3’ tRFs in the analyzed probes respectively (Figure 2B). Interestingly, we supports the idea proposed by Schorn and coworkers ascertained a substantial difference considering the who assume that this mechanism could be highly abundance of tRNA fragments (tRFs). In both probes, conserved across different species, providing an 21 nt RNAs derived from the 3’ end of tRNAs (3’ tRFs) innate immunity against LTR propagation. constitute the vast majority of tRNA fragments.

**Figure 2.**
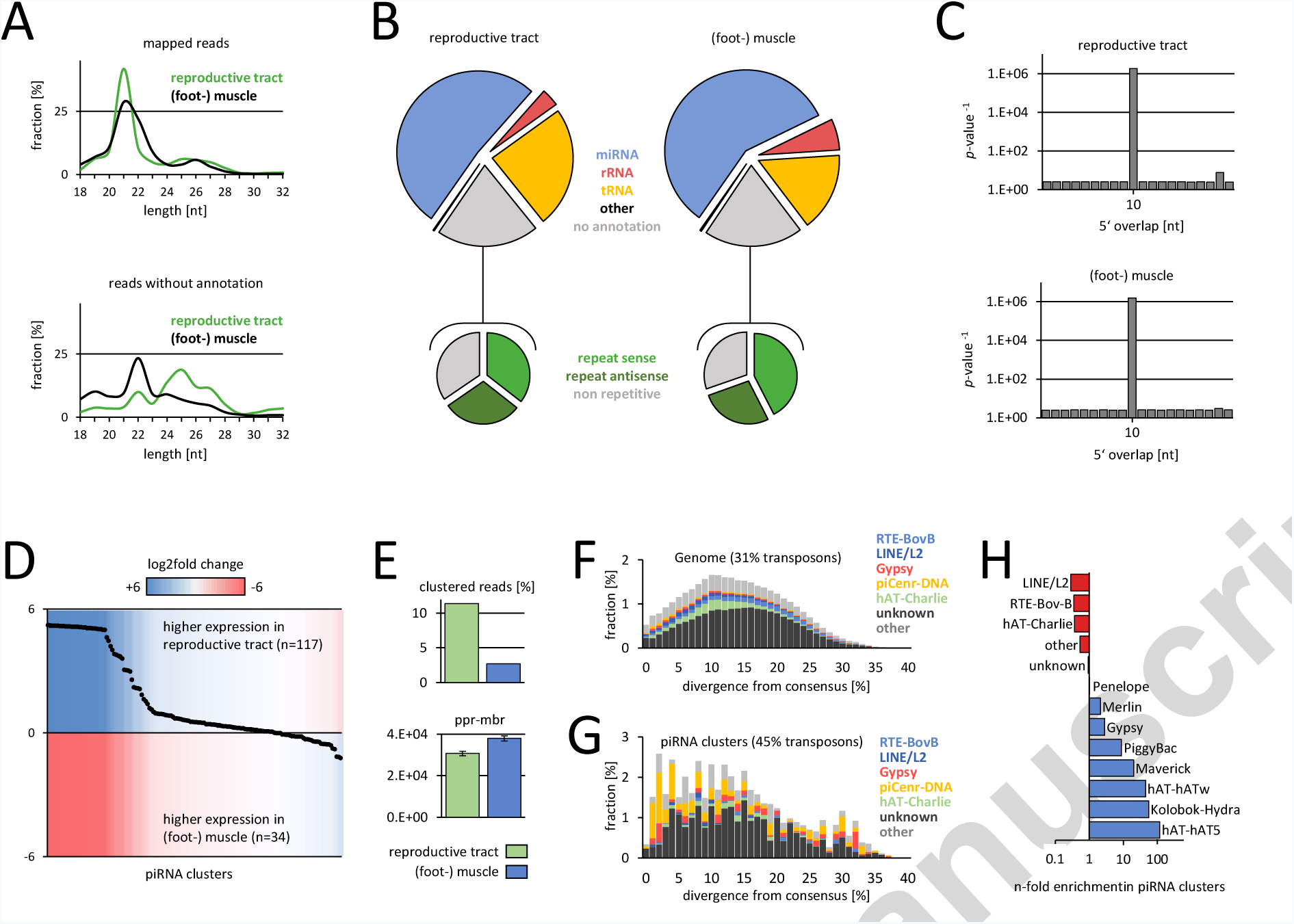
Characterization of small RNAs from*L. stagnalis*(foot-) muscle and reproductive tract. (A) Sequence read length distribution of mapped(top) and unannotated reads (bottom). (B) Small RNA annotation (top) and transposon content of unannotated reads (bottom). (C) Ping-pong signature. P-values are deduced from the corresponding Z-scores. (D) Differential expression of 151 predicted piRNA clusters. (E) Amount of clustered reads (top) and ping-pong reads per million bootstrapped reads (bottom). (F) Representation of transposons in the genome of *L.stagnalis*, plotted by divergence [%] from transposon consensus. (G) Representation of transposons within piRNA clusters of *L. stagnalis*, plotted by divergence [%] from transposon consensus. (H) Transposons that are enriched or depleted in *L. stagnalis* piRNA clusters.

Focusing on putative piRNAs, we analyzed the fraction of sequence reads that did not match to any other class of non-coding RNA. This dark matter of sRNAs comprises 18% and 17% of sequence reads in the reproductive tract and in muscle, respectively.

Strikingly, these sRNAs are strongly enriched for transposon sequences, suggesting their involvement in transposon control (Figure 2B). To verify the presence of piRNAs, we checked for the so-called ping-pong signature (bias for 10 bp 5’ overlap of mapped sequence reads), which is a hallmark of secondary piRNA biogenesis and requires the catalytic activity - and thus expression - of PIWI proteins (Czech and Hannon 2016). Remarkably, we detected a significant ping-pong signature in both, the reproductive tract and muscle (Figure 2C), suggesting active PIWI/piRNA-dependent transposon silencing in the germline and the soma. Next, we used proTRAC (Rosenkranz and Zischler 2012) to identify 151 piRNA clusters, covering 0.13% of the genome of *L. stagnalis* (Figure 2D). Although piRNAs originate from essentially identical clusters in the reproductive tract and in muscle, we found that 11.4% of sequence reads from the reproductive tract map to piRNA clusters, while only 2.7% of sequence reads from muscle do so, indicating rather moderate production of primary piRNAs in the soma compared to the germline (Figure 2E). Nevertheless, we found that the number of piRNAs that participate in ping-pong-amplification (ping-pong reads per million bootstrapped reads, ppr-mbr) is even slightly higher in muscle compared to the situation in the reproductive tract, emphasizing the functional importance of somatic PIWI/piRNA expression (Figure 2E). In line with the transposon-suppressive role of piRNAs, the identified piRNA clusters are clearly enriched for transposon sequences compared to the whole genome situation (45% and 31%, respectively, figure 2F and 2G). Interestingly, the transposon composition in piRNA clusters does not reflect the transposon landscape of the genome. Instead, piRNA clusters are enriched for Gypsy retrotransposons and particularly DNA transposons such as PiggyBac or hAT5, showing up to 120-fold enrichment in piRNA clusters (figure 2G and 2H). This non-random distribution suggests a selective regime that favors insertion events of evolutionary young and active transposons.

### Ubiquitous and dynamic expression of piRNAs in *Crassostrea gigas*

Based on our observation that PIWI genes and piRNAs are expressed in the soma and the germline of *L. stagnalis*, we reanalyzed previously published small RNA datasets from *C. gigas* that were used to investigate the dynamic expression of miRNAs during oyster development without further examination of a putative piRNA fraction (Xu et al. 2014, NCBI Sequence Read Archive Project ID SRP007591). We annotated *C. gigas* sRNAs from the male and female gonad, different developmental stages ranging from the egg to juvenile, and a representative set of somatic tissues from adult animals (Supplementary Table 1). In all datasets, particularly in gonadal tissues, eggs and early embryo stages but also in hemolymph we detected a large amount of sequence reads that did not match to any known ncRNA class but was instead enriched for transposon sequences. The transposon-matching sub-fraction itself was enriched for antisense sequences (Supplementary Table 1). Analogous to the procedure applied for the *L. stagnalis* datasets, we verified the presence of primary and secondary piRNAs by analyzing the ping-pong signature of each dataset. Remarkably, we detected a significant ping-pong signature across all analyzed datasets (Figure 3A, Supplementary Figure 1), but also found that the number of ping-pong reads (measured as ppr-mbr) differs considerably depending on the tissue and developmental stage (Figure 3A, Supplementary Figure 2). Noteworthy, a ping-pong signature is also detectable when taking only those reads into account that match protein coding sequences, suggesting a relevant role of the PIWI/piRNA pathway in post-transcriptional regulation of protein coding genes in gonads, egg blastula, digestive gland and hemolymph (Supplementary Table 2). We further used sequences without ncRNA annotation to predict piRNA clusters with proTRAC and checked whether we can observe a differential expression of specific piRNA clusters in time and space (Figure 3A).

In contrast to the situation in *L. stagnalis*, we found that different genomic loci are responsible for production of primary piRNAs in the germline and in the soma, but also during different developmental stages. A clustering approach based on average linkage (Babicki et al. 2016) revealed four distinct groups of piRNA clusters which we named class 1-4 piRNA clusters (Figure 4A). Class 1 piRNA clusters are active in the adult germline (male and female) and in the early embryo until the D-shaped veliger stadium where larvae are approximately 14 hours old. The same applies to class 2 piRNA clusters, however, following the D-shape veliger stadium, class 1 piRNA clusters become inactive, while class 2 piRNA clusters remain active and class 3 piRNA clusters start piRNA production. Both, class 2 and class 3 piRNA cluster activity is measurable until the juvenile stadium, where oysters are approximately 20 days old. In somatic tissues of adult oysters, class 4 piRNA clusters represent the main source of primary piRNAs (Figure 4A, bottom). Interestingly, all four classes of piRNA clusters are active in hemocytes, which also feature the highest amount of clustered reads, and ping-pong reads compared to other somatic tissues. This might reflect the presence of stem cells within the hemocyte cell population which is subject to complex differentiation processes (Fisher 1986, Lau et al. 2017).

**Figure 3.**
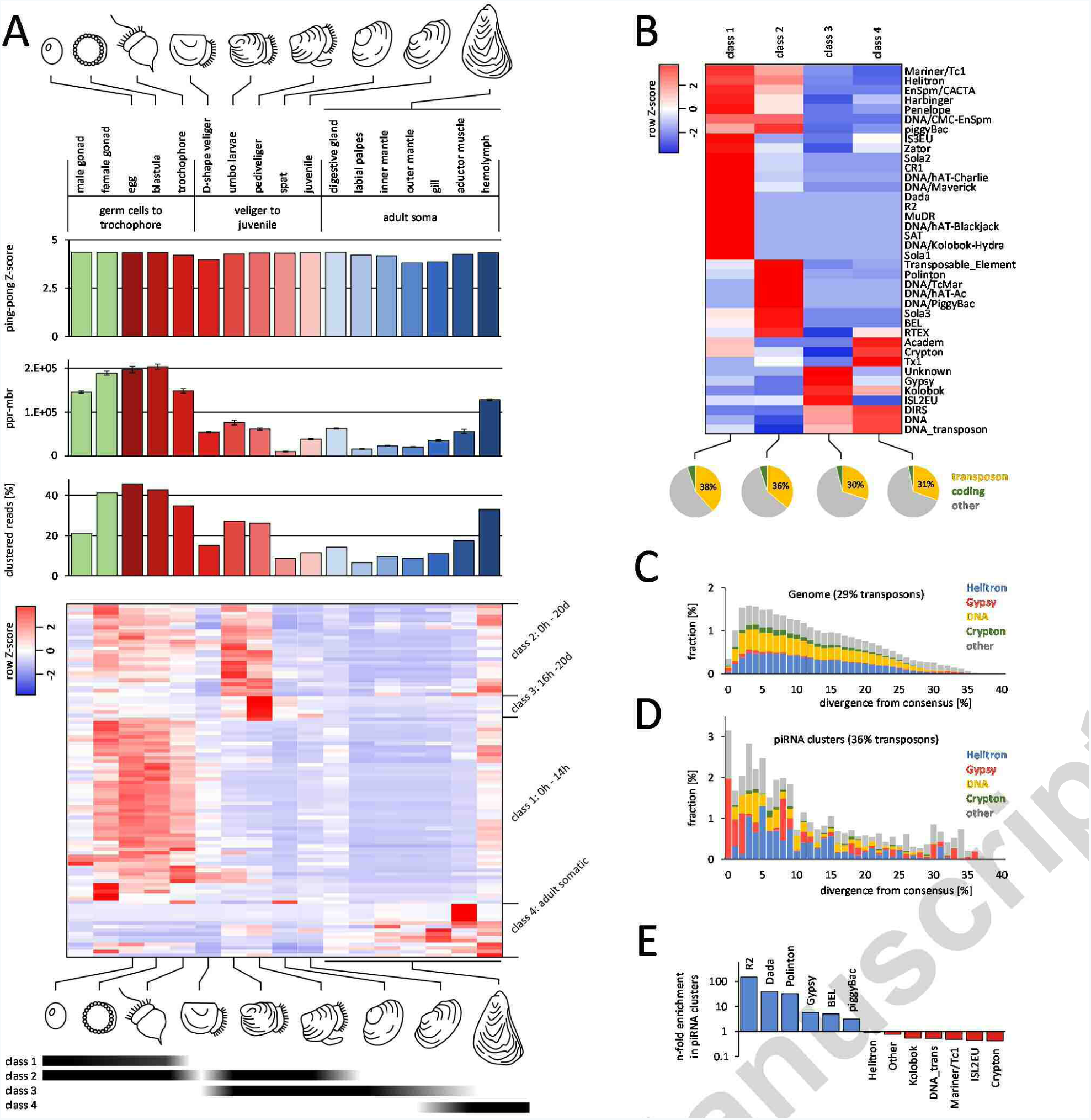
Characterization of small RNAs from different *C. gigas*samples. (A) Sequence reads without annotation produce a significantping-pong signature (top row of bars, only Z-scores for 10 bp 5’ overlap are shown). The number of ping-pong reads per million bootstrapped reads (middle row of bars), and the number of clustered reads (bottom row of bars) differs considerably across the probes. Heatmap shows the differential expression of the top 100 piRNA clusters in terms of maximum rpm coverage. Different classes of piRNA clusters are expressed during oyster development and in adult somatic tissues (bottom). (B) Transposon composition of piRNA clusters belonging to four different classes. (C) Representation of transposons in the genome of *C. gigas*, plotted by divergence [%] from transposon consensus. (D) Representation of transposons within piRNA clusters of *C. gigas*, plotted by divergence [%] from transposon consensus. (E) Transposons that are enriched or depleted in *C. gigas* piRNA clusters.

Interestingly, the four classes of piRNA clusters differ generally enriched for transposon sequences showing considerably regarding the overall transposon 38% and 36% transposon derived sequences, content as well as the specific transposon composition respectively, compared to a genomic transposon (Figure 3B-D). Class 1 and class 2 piRNA clusters are content of 29%. The surprisingly high accumulation of young (as deduced from the divergence from their consensus) Gypsy elements in piRNA clusters, suggests a strong selection for Gypsy element insertions, probably as a consequence of Gypsy activity in *C. gigas*. Considering transposons that are generally enriched in piRNA clusters (Figure 3E) we found that R2 retrotransposons (149-fold enrichment in piRNA clusters) and Dada DNA transposons (40-fold enrichment in piRNA clusters) are most abundant in class 1 piRNA clusters. In contrast, Polinton DNA transposons (32-fold enrichment in piRNA clusters) and BEL retrotransposons (5-fold enrichment in piRNA clusters) are most abundant in class 2 piRNA clusters. Different from class 1 and class 2 piRNA clusters, class 3 and class 4 piRNA clusters display only slight transposon enrichment (30% and 31%, respectively). Noteworthy, high copy number Gypsy retrotransposons (5-fold enrichment in piRNA clusters) are most abundant in class 3 piRNA clusters, while Academ, Crypton and Tx1 transposons are most abundant in class 4 piRNA clusters.

These results contrast with the situation in *L. stagnalis*, where identical piRNA producing loci are active in the germline and in the soma. Moreover, we can observe considerable differences in the transposon composition of piRNA clusters in the two species, which likely reflect divergent transposon activity and resulting selective constraints on the different phylogenetic lineages.

### Homotypic and heterotypic ping-pong amplification

The so-called ping-pong amplification loop describes a process that is responsible for the post-transcriptional silencing of transposable elements. In *Drosophila* and mouse, this process typically involves two PIWI paralogs (heterotypic ping-pong), one loaded with antisense piRNAs targeting transposon transcripts, and the other loaded with sense piRNAs targeting piRNA cluster transcripts, which contain transposon sequences in antisense orientation (Brennecke et al. 2007, Aravin et al. 2008). Likely for steric reasons, premature piRNAs loaded onto the different PIWI paralogs are more or less rigorously trimmed at their 3’ ends. This is why piRNA populations bound to different PIWI paralogs not only differ regarding the amount of sense- and antisense-transposon sequences, but also in their sequence length profiles (Aravin et al. 2007, Kawaoka et al. 2011, Czech and Hannon 2016). In addition to the heterotypic ping-pong amplification, homotypic ping-pong has been shown to occur in *qin* mutant flies (Aub:Aub, Zhang et al. 2011), and wildtype prenatal mouse testis (Miwi2:Miwi2, Mili:Mili, Aravin et al. 2008).

Since the typical molluskan genome encodes two ubiquitously expressed PIWI paralogs, Piwil1 and Piwil2, we asked whether we are able to show the participation of distinct piRNA populations in the ping-pong cycle. We conducted a bioinformatics approach under the premise that Piwil1- and Piwil2-bound piRNAs exhibit different length profiles, which is the case for the corresponding mouse homologs Piwil1 (Miwi) which preferentially binds 29/30 nt piRNAs, and Piwil2 (Mili), which preferentially binds 26/27 nt piRNAs (Vourekas et al. 2012). We analyzed pairs of mapped *C. giga*s and *L. stagnalis* sequence reads that showed a 10 bp 5’ overlap (ping-pong pairs), with respect to the sequence length of each ping-pong partner (Figure 4, Supplementary Figure 3). In the female gonad of *C. gigas*, most ping-pong pairs combine piRNAs with a length of 25 nt and 29 nt, which suggests heterotypic ping-pong amplification. (Figure 4) In support of this, 29 nt piRNAs, presumably bound to Piwil1, are heavily biased for a 5’ uridine (a hallmark of primary piRNAs), whereas 25 nt piRNAs, presumably bound to Piwil2, show a stronger bias for an adenine at position 10 (typical for secondary piRNAs). In contrast, ping-pong pairs in *C. gigas*muscle predominantly combine two 29 nt piRNAs, suggesting homotypic, Piwil1-based ping-pong amplification. Generally, the observed patterns of ping-pong pairs are very diverse across the different samples, for instance displaying heterotypic ping-pong in the digestive gland and homotypic, Piwil2-based ping-pong in hemolymph cells (Supplementary Figure 3).

Since the expression of Piwil1 compared to Piwil2 is considerably lower in *L. stagnalis*, we were curious to check whether the corresponding ping-pong pairs might reflect this fact. Indeed, 26/26 nt pairs (homotypic, Piwil2-based ping-pong) represent the majority of ping-pong pairs in the reproductive tract, followed by 25/28 nt pairs (Figure 4). In addition, homotypic Piwil2-based ping-pong amplification with 24/25 nt ping-pong pairs is also dominant in the *L. stagnalis* muscle.

## Conclusions

Our results reveal that mollusks utilize the PIWI/piRNA pathway as a defense against transposable elements in the germline and in the soma, which corresponds to the situation in arthropods and therefor suggests somatic PIWI/piRNA expression as an ancestral bilaterian character state. In addition, based on the observation that a substantial fraction of arthropod and oyster piRNAs targets messenger RNAs (a gene annotation for *L. stagnalis* was not available) it seems likely that the last common ancestor of arthropods and mollusks applied the PIWI/piRNA pathway also for post-Somatic PIWI/piRNA expression in mollusks transcriptional regulation of protein coding genes. In vertebrates, somatic PIWI/piRNA expression appears to have fade away and reports on somatically expressed piRNAs in mammals are often considered with skepticism. However, remnants of the former somatic expression might have outlasted to fulfill special functions in specific cells and/or in narrowly defined timespans of development or cell differentiation in the one or the other clade. In any case, we should be aware of the attitude that experiments with *Drosophila* and mouse will tell us everything that is worth to know about the PIWI/piRNA pathway.

**Figure 4.**
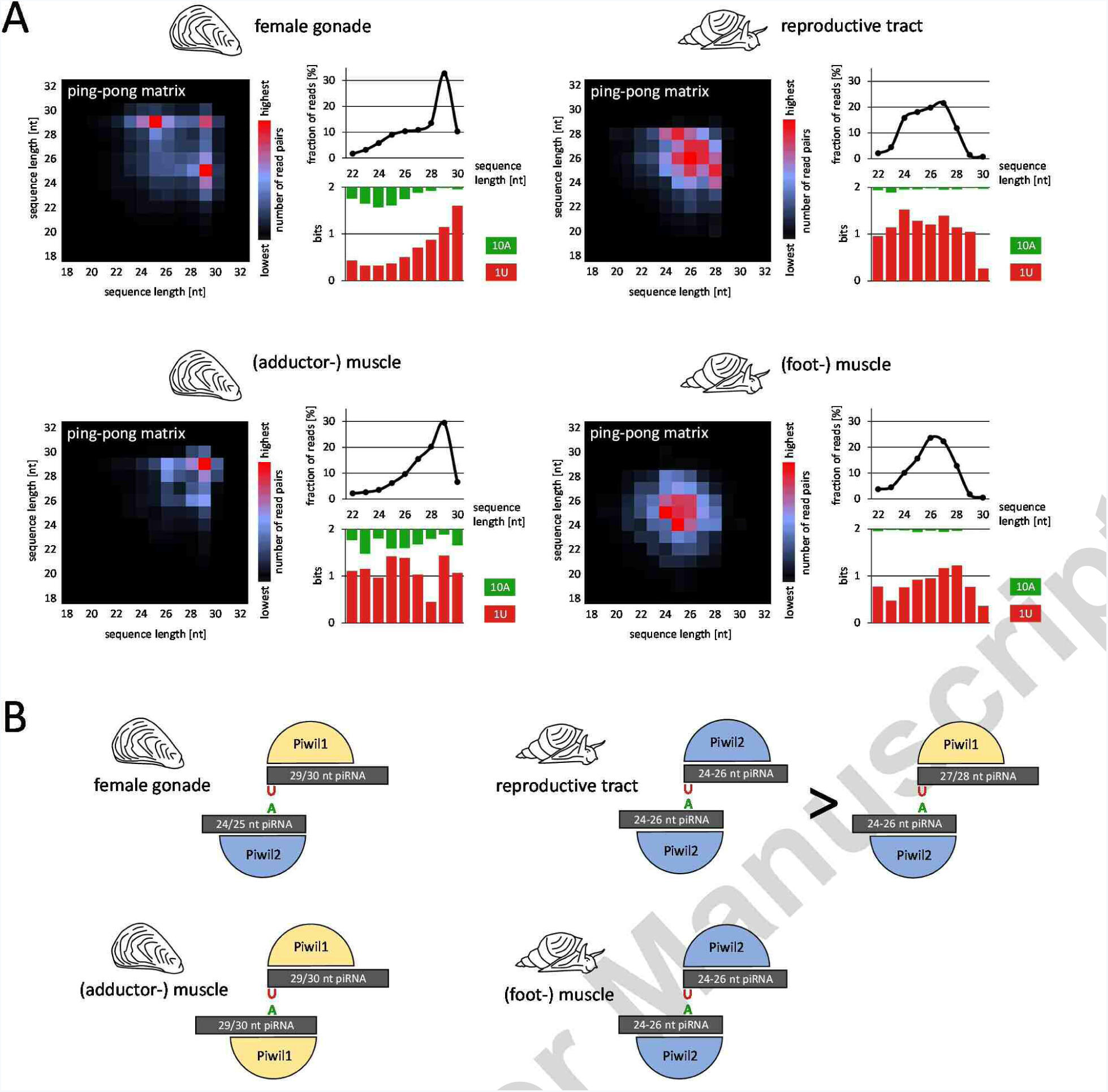
Analysis of piRNAs that participate in the ping-pong amplification loop. (A) Ping-pong matrices illustrate frequent length-combinations of ping-pong pairs (sequences with 10 bp 5’ overlap). Sequence read length distribution and 1U/10A bias [bits] for ping-pong sequences are shown. (B) Proposed model of ping-pong amplification in the germline and muscle of *C. gigas* and *L. stagnalis*.

## Material and Methods

### Piwi gene annotation and tree reconstruction

In order to reconstruct the phylogenetic relations of mollusk Piwi proteins, we first searched for Piwi genes in species with an available genome sequence that lack proper annotation (*Lymnaea stagnalis*, *Radix auricularia*, *Lottia gigantea*, *Bathymodiolus platifrons*, *Pinctada martensii*). To this end, we scanned the relevant genomes for sequences that are homologous to annotated Piwi paralogs of the pacific oyster (EKC35279 and EKC29295) by aligning translated DNA sequences using tblastx. Neighboring hits with a distance smaller than 10 kb were grouped as exons of distinct gene loci. Only groups containing the overall best hits for a given locus were retained. Finally, the predicted gene homologs were checked for presence of PIWI and PAZ domains using NCBI conserved domain database (Marchler-Bauer et al. 2015). Similarly, for Piwi expression analysis by qPCR in the pond snail, we identified the housekeeping gene GPI (glucose-6-phosphate isomerase) by comparison with the human ortholog (ARJ36701). The predicted and annotated Piwi protein sequences of all available molluskan species together with human Piwi paralogs (Piwil1-4) and Drosophila Ago3 were aligned using MUSCLE (Edgar 2004). The resulting protein alignment was then used for phylogenetic tree reconstruction with PhyML (Guindon et al. 2009) using approximate likelihood-ratio test (SH-like) and Dayhoff substitution model.

### qPCR

To estimate the expression of the Piwil homologs in several tissues of *L. stagnalis* and *C. gigas* we performed qPCR with cDNA synthesized from the total RNA fraction of these tissues. Total RNA was isolated with TriReagent and the polyadenylated transcriptome was reversely transcribed with SuperScript IV using the RT-primer 5’-CGAATTCTAGAGCTCGAGGCAGGCGACATGT25VN-3’. Primers amplifying ~ 200 bp long products of the respective Piwil homologs and housekeeping genes were designed with the NCBI tool primer-BLAST on basis of the *L. stagnalis* genome scaffold (14639) and the *C. gigas* genome assembly (10758). To prevent amplification of residual genomic DNA, primers were designed to be exon-junction spanning or to span at least several intronic regions. The respective biological replicates were analyzed as technical duplicates on a Corbett Rotor-Gene 6000 real-time PCR cycler and the transcript expression was quantified by standard curves of the individual primer pair amplicons. For each cDNA sample the calculated PIWI concentrations were finally normalized by the calculated copy numbers of the housekeeping genes.

### Small RNA extraction and sequencing

Experiments were performed on commercially available *C. gigas* animals from the western French Atlantic coast (lle d’Oleron) and captured wild living *L. stagnalis* animals from South-western Germany (Heppenheim). We extracted total RNA from *L. stagnalis* reproduction tract (incl. ovotestis, oviduct, spermatheca, spermiduct, prostate, uterus, vagina, vas deferenc) and foot muscle, and total RNA from *C. gigas* adductor muscle and gonadal tissue with TriReagent according to the manufacturer’s instructions. For each species we sampled two different individuals per tissue. The small RNA fractions of each obtained total RNA probe were sequenced at BGI, Hong Kong, on a BGISEQ-500 unit. Sequence datasets are deposited at NCBI’s Sequence Read Archive (SRA) and can be accessed under the SRA project ID SRP123456. We further used previously published small RNA sequence data from *C. gigas* (Xu et al. 2014) to analyze piRNA expression and characteristics with respect to different developmental stages.

### Repeat annotation

We performed *de novo* prediction of repetitive elements in the genome of *L. stagnalis* with RepeatScout (v. 1.0.5, Price et al. 2005). Predicted repetitive elements were classified with Repeat Classifier which is part of the RepeatModeler (v. 1.0.11) package. The resulting repeat sequences, as well as a complete collection of currently available molluskan repeat sequences from RepBase (Bao et al. 2015) were used as reference sequences for repeat masking of the *L. stagnalis* and *C. gigas* genomes with Repeat Masker (v. 4.0.7) using the cross_match search engine and the option -s for most sensitive masking.

### Processing and annotation of small RNA sequence data

Small RNA sequence datasets were collapsed to non-identical sequences, retaining information on sequence read counts using the Perl script *collapse*. Sequences >36nt were rejected using the Perl script *length-filter*. Finally, low complexity sequences were filtered using the Perl script *duster* with default parameters. All Perl scripts mentioned are part of the NGS toolbox (Rosenkranz et al. 2015).

We then applied a customized mapping strategy of the remaining small RNA sequence reads based on the consideration that our datasets presumably contain considerable amounts of transposon- derived piRNAs as well as post-transcriptionally edited (e.g. A-to-I) or tailed miRNAs and piRNAs. Genomic mapping was performed with SeqMap (Jiang and Wong 2008) using the option /output_all_matches and allowing up to three mismatches. The obtained alignments were further filtered using an in-house Perl script that is available upon request. For the final alignments we allowed up to two non-template 3’ nucleotides and up to one internal mismatch. For each sequence, we only considered the best alignments in terms of mismatch counts, but did not reject alignments with equal quality in case of multiple mapping sequences. Sequences that did not produce at least one valid alignment to the reference genome were rejected.

To improve small RNA sequence annotation, we performed *de novo* tRNA, rRNA and miRNA prediction based on the available reference genome assemblies GCA_900036025.1 v1.0 (*L. stagnalis*) and GCA_000297895.1 oyster_v9 (*C. gigas*). tRNA annotation was performed with a local copy of tRNAscan (v. 1.3.1, Lowe and Chan 2016). Only tRNAs with less than 5% N’s were taken for further analysis. rRNA sequences were predicted using a local copy of RNAmmer (v. 1.2, Lagesen et al. 2007) and hmmer (v. 2.2g, Johnson et al. 2010). Both tools were run with default parameters. We pooled small RNA sequence reads from different replicates and tissues for each species separately to perform miRNA *de novo* prediction with ShortStack (v. 3.8.4, Axtell 2013) using default parameters. The predicted tRNA, rRNA and miRNA precursor sequences, as well as previously published miRNA precursor sequences (Xu et al. 2014, Zhou et al. 2014, Zhao et al. 2016), were used as additional reference sequences for small non-coding RNA annotation with unitas (v.1.4.6, Gebert at. al. 2017).

### piRNA cluster identification

Sequences that did not produce a match to known non-coding RNAs were considered as putative piRNAs and were used for prediction of piRNA clusters with proTRAC (v. 2.4.0) applying default settings. piRNA clusters were predicted for each dataset and species separately. The resulting piRNA cluster predictions for each species were condensed, merging clusters with less than 10 kb distance from each other. Finally, we calculated the sequence read coverage [rpm] for each of the resulting piRNA clusters per dataset. For *C. gigas* piRNA clusters, a heat map for the top 100 piRNA clusters in terms of maximum rpm coverage (accounting for 64% of summed rpm values) was constructed with Heatmapper (Babicki et al. 2016) applying Pearson distance and average linkage clustering. For *L. stagnalis* piRNA clusters we

### Ping-pong quantification

In order to compare ping-pong signatures across multiple datasets with different sequencing depth, we constructed a software tool, PPmeter, that creates bootstrap pseudo-replicates from original datasets and subsequently analyzes the ping-pong signature and number of ping-pong sequence reads of each pseudo-replicate (default: 100 pseudo-replicates each comprising one million sequence reads). The obtained parameters ‘ping-pong score per million bootstrapped reads’ (pps-mbr) and ‘ping-pong reads per million bootstrapped reads’ (ppr-mbr) can be used for quantification and direct comparison of ping-pong activity in different small RNA datasets. The software is freely available at http://www.smallRNAgroup.uni-mainz.de/software.html.

## Supplementary Material

**Supplementary Table 1.**
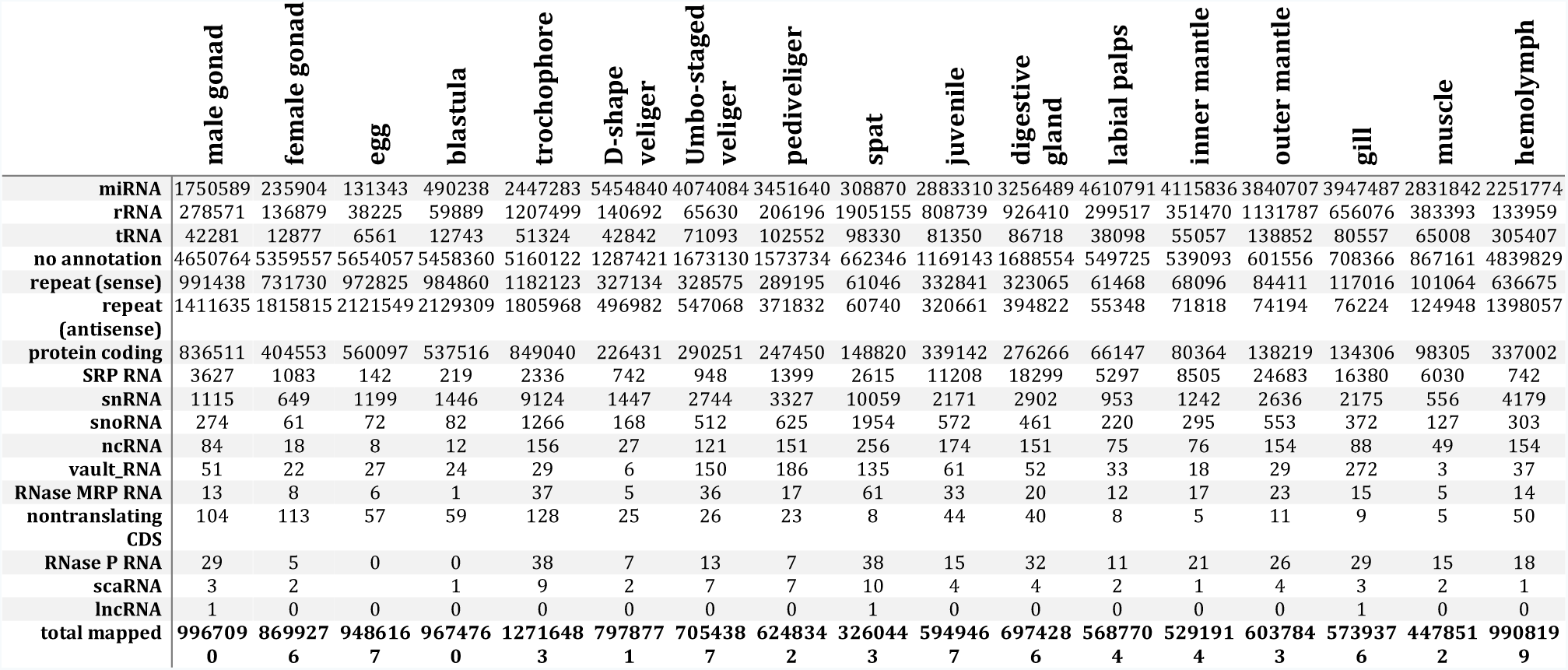
Annotation of small RNAs from *C.gigas* samples. Reads without annotation or reads that match transposon (repeat) sequences represent putative piRNAs. Read counts of multiple mapping sequences were fractionated accordingly. Values are rounded, which explains possible discrepancies with the total number of mapped reads.

**Supplementary Table 2.** Counts of sequence read pairs with a specific 5’ overlap considering only those sequences that match protein coding sequences.

| 5' overlap | male gonad | female gonad | egg | blastula | trochophore | D-shape veliger | Umbo-staged veliger | pediveliger | spat | juvenile | digestive gland | labial palps | inner mantle | outer mantle | gill | muscle | hemolymph |
| --- | --- | --- | --- | --- | --- | --- | --- | --- | --- | --- | --- | --- | --- | --- | --- | --- | --- |
| 1 | 741 | 201 | 12711 | 58374 | 19108 | 15174 | 350038 | 2965 | 0 | 377 | 279 | 2 | 1 | 0 | 1 | 3 | 139 |
| 2 | 364 | 525 | 571 | 1172 | 452 | 61 | 979 | 39 | 0 | 98 | 418 | 1 | 0 | 0 | 0 | 0 | 366 |
| 3 | 538 | 263 | 843 | 1348 | 1992 | 561 | 43179 | 1321 | 1 | 80 | 273 | 6 | 1 | 5 | 1 | 14 | 73 |
| 4 | 927 | 454 | 1584 | 2955 | 3190 | 3989 | 15734 | 365 | 1 | 60 | 257 | 0 | 0 | 2 | 1 | 0 | 101 |
| 5 | 1302 | 577 | 2047 | 1751 | 1234 | 104 | 7181 | 156 | 0 | 212 | 279 | 2 | 1 | 0 | 0 | 1 | 112 |
| 6 | 398 | 668 | 726 | 759 | 668 | 219 | 2026 | 184 | 1 | 84 | 306 | 1 | 0 | 0 | 0 | 3 | 90 |
| 7 | 1962 | 3310 | 12088 | 6555 | 10252 | 497 | 961 | 74 | 0 | 92 | 753 | 0 | 1 | 1 | 1 | 0 | 1604 |
| 8 | 2161 | 670 | 2111 | 2474 | 2301 | 635 | 4496 | 87 | 1 | 222 | 244 | 0 | 1 | 0 | 0 | 2 | 442 |
| 9 | 2128 | 755 | 2278 | 2421 | 977 | 597 | 418 | 10 | 3 | 76 | 478 | 0 | 0 | 1 | 0 | 0 | 10169 |
| 10 | 65665 | 22281 | 44756 | 57158 | 26768 | 7908 | 4253 | 378 | 5 | 4536 | 50170 | 22 | 10 | 16 | 24 | 9 | 167909 |
| 11 | 1355 | 687 | 1409 | 1263 | 1738 | 153 | 72 | 9 | 0 | 25 | 268 | 0 | 0 | 0 | 1 | 2 | 279 |
| 12 | 1262 | 634 | 1730 | 7732 | 1299 | 1418 | 816 | 154 | 1 | 257 | 297 | 0 | 0 | 1 | 0 | 4 | 624 |
| 13 | 900 | 5312 | 2765 | 2857 | 5098 | 520 | 18017 | 633 | 1 | 85 | 4646 | 2 | 5 | 11 | 2 | 1 | 9229 |
| 14 | 5589 | 3337 | 5842 | 19404 | 10693 | 13444 | 2184 | 758 | 1 | 150 | 59 | 0 | 0 | 0 | 0 | 1 | 58 |
| 15 | 2964 | 1386 | 2208 | 5560 | 2747 | 3068 | 2869 | 529 | 1 | 215 | 58 | 0 | 0 | 0 | 1 | 3 | 92 |
| 16 | 2471 | 589 | 723 | 676 | 466 | 91 | 278 | 65 | 8 | 30 | 66 | 1 | 0 | 6 | 1 | 29 | 174 |
| 17 | 418 | 345 | 419 | 432 | 281 | 85 | 45 | 81 | 9 | 33 | 19 | 0 | 0 | 1 | 0 | 0 | 27 |
| 18 | 979 | 1000 | 760 | 577 | 1207 | 143 | 131 | 36 | 12 | 41 | 24 | 1 | 0 | 0 | 1 | 1 | 63 |
| 19 | 962 | 262 | 250 | 298 | 152 | 61 | 84 | 19 | 7 | 13 | 27 | 8 | 3 | 17 | 115 | 1 | 162 |
| 20 | 1398 | 310 | 664 | 591 | 378 | 160 | 184 | 32 | 10 | 73 | 23 | 0 | 0 | 0 | 0 | 0 | 35 |

**Supplementary Figure 1.**
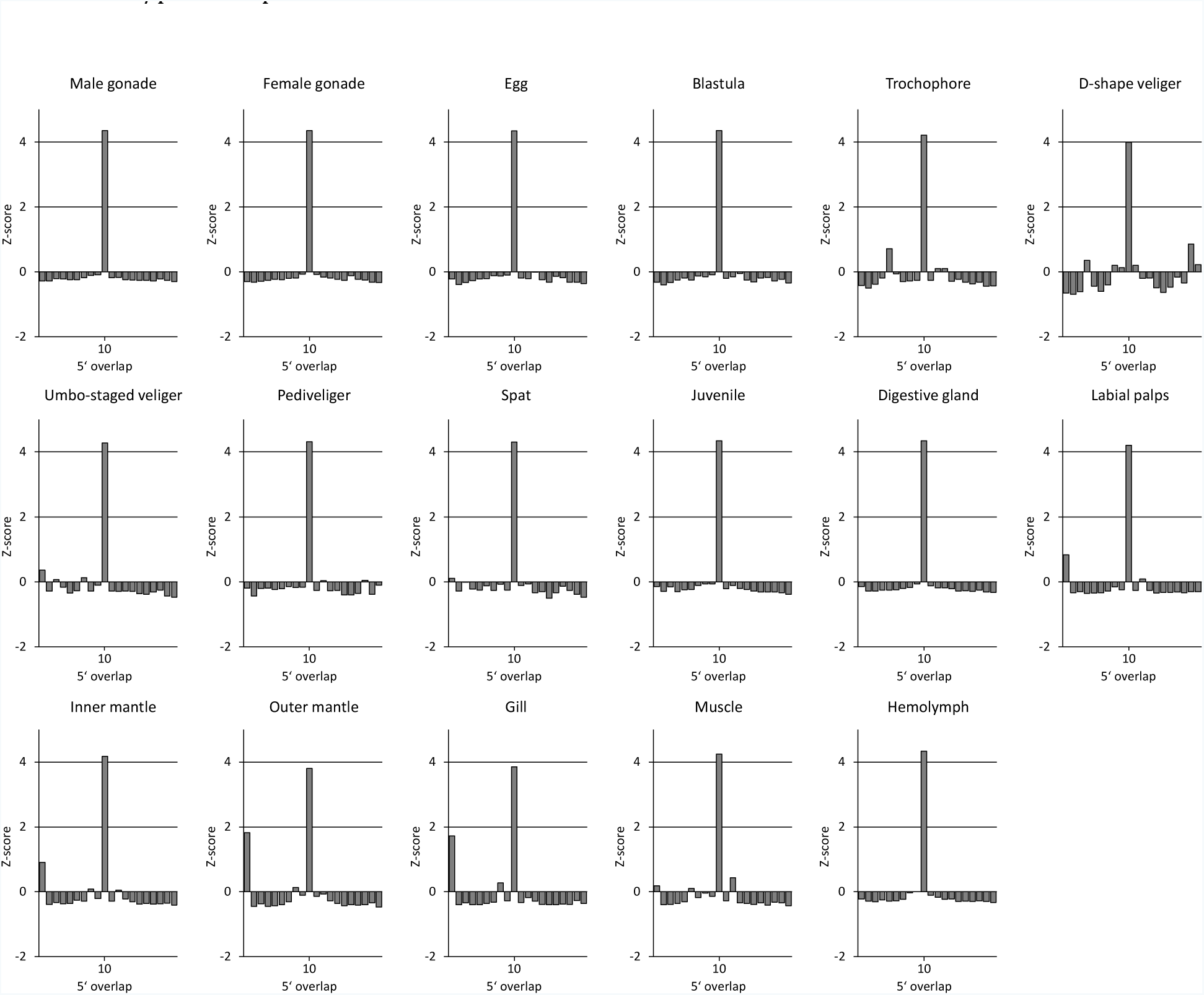
Ping-pong signature of small RNAs from *C. gigas* samples.

**Supplementary Figure 2.**
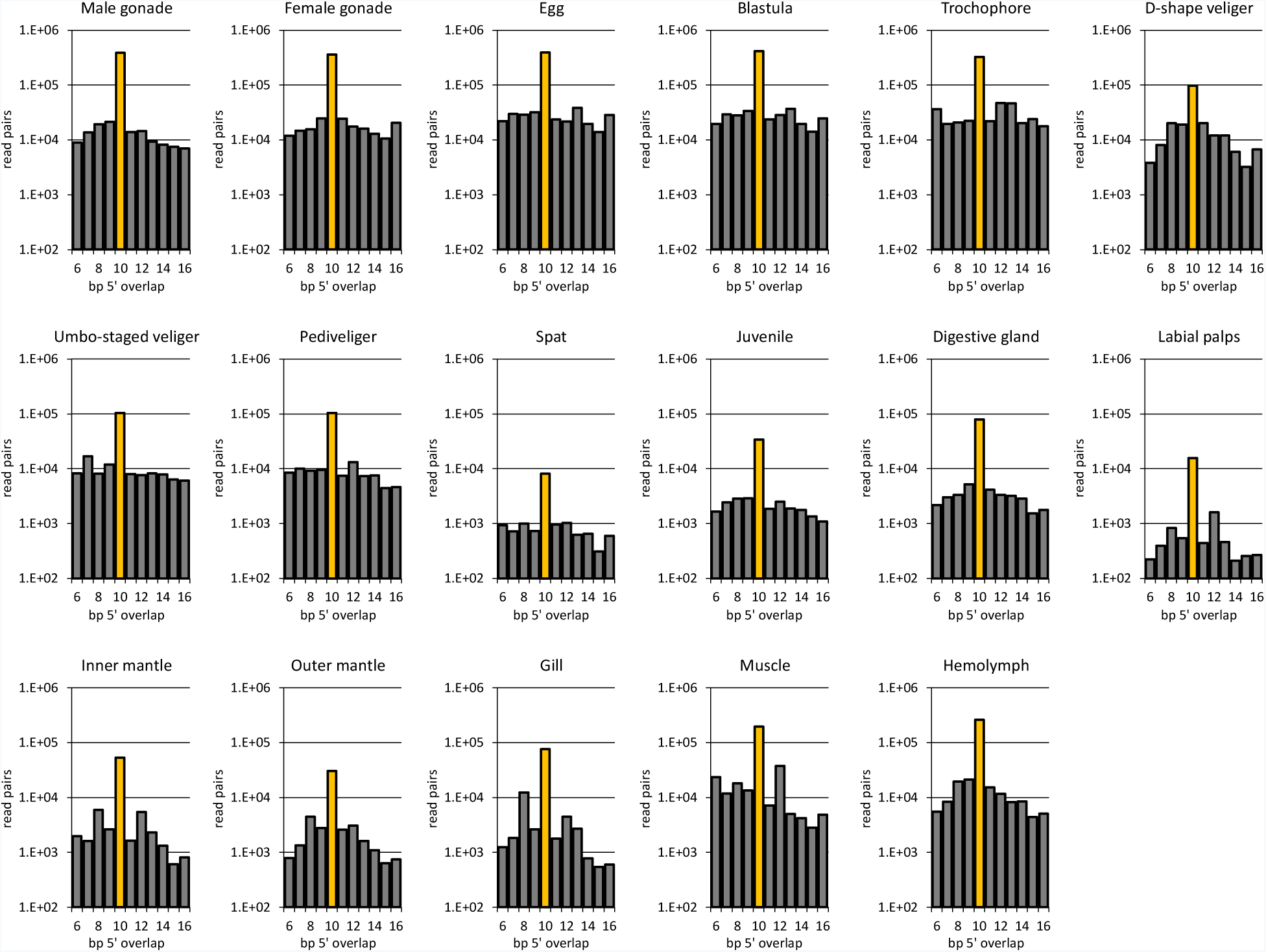
Graphs depict the average number of sequence read pairs with a specific 5‘ overlap for 100 pseudo-replicates (PR) per dataset with one million reads per PR. The value for read pairs with 10 nt overlap can serve as a measurement for the intensity of ping-pong amplification and is also directly comparable across different datasets.

**Supplementary Figure 3.**
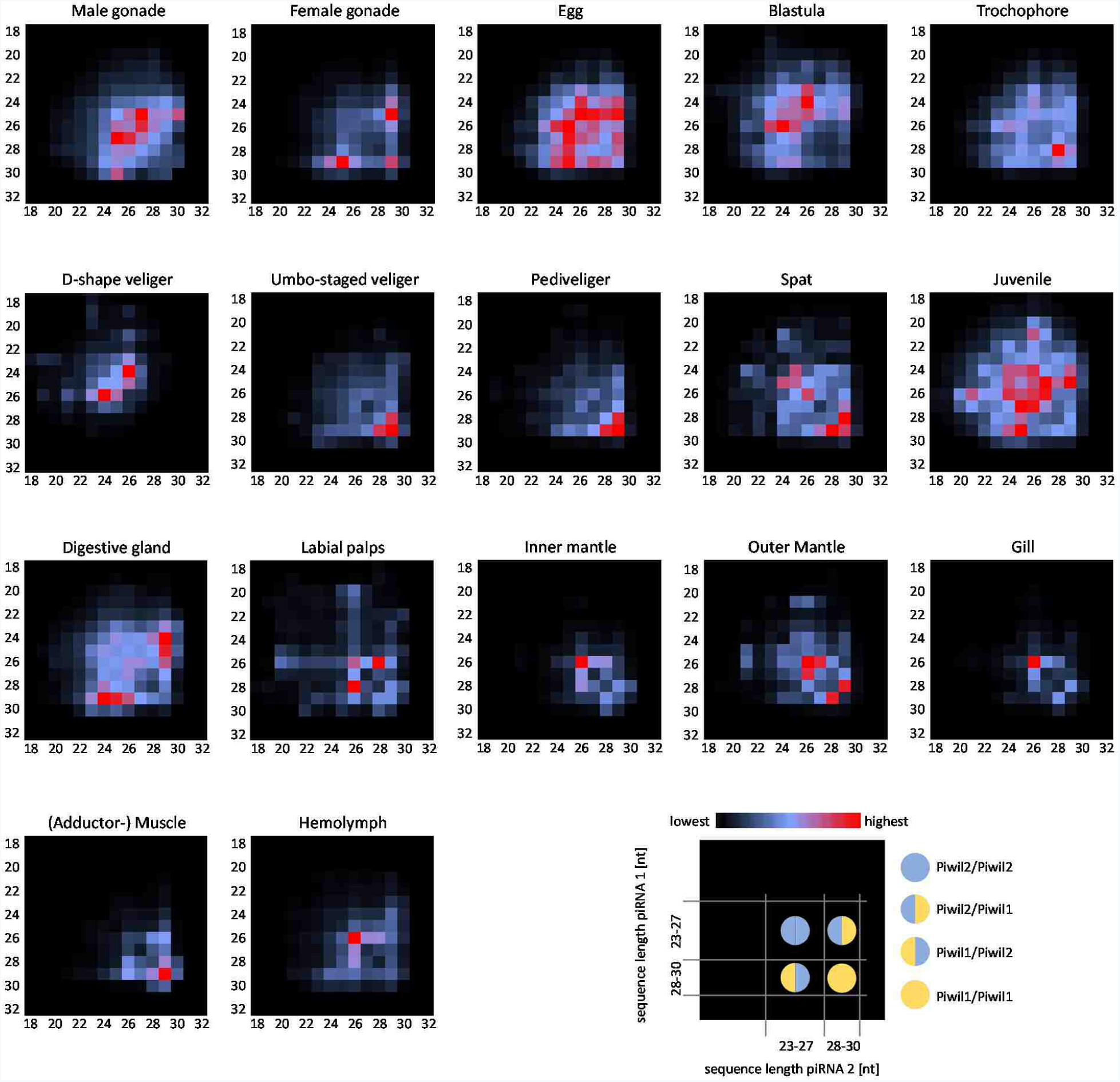
Ping-pong matrices for small RNA samples from *C. gigas*. Frequent length-combinations of ping-pong pairs (sequences with 10 bp 5’ overlap) are indicated in red. x-axis and y-axis refer to sequence read length of the two sequences of a ping-pong pair.

## References

Aravin AA, Hannon GJ, Brennecke J., The Piwi-piRNA pathway provides an adaptive defense in the transposon arms race. Science. 2007, 318(5851):761–4.

Aravin AA, Sachidanandam R, Bourc'his D, Schaefer C, Pezic D, Toth KF, Bestor T, Hannon GJ. A piRNA pathway primed by individual transposons is linked to de novo DNA methylation in mice. Mol Cell. 2008, 31(6):785–99.

Axtell MJ. ShortStack: comprehensive annotation and quantification of small RNA genes. RNA. 2013, 19(6):740–51.

Babicki S, Arndt D, Marcu A, Liang Y, Grant JR, Maciejewski A, Wishart DS. Heatmapper: web-enabled heat mapping for all. Nucleic Acids Res. 2016, 44(W1):W147–53.

Bao W, Kojima KK, Kohany O., Repbase Update, a database of repetitive elements in eukaryotic genomes. Mob DNA. 2015, 6:11.

Brennecke J1, Aravin AA, Stark A, Dus M, Kellis M, Sachidanandam R, Hannon GJ., Discrete small RNA-generating loci as master regulators of transposon activity in Drosophila. Cell. 2007, 128(6):1089–103.

Czech B, Hannon GJ., One Loop to Rule Them All: The Ping-Pong Cycle and piRNA-Guided Silencing. Trends Biochem Sci. 2016, 41(4):324–37.

Giacomo M1 Di, Comazzetto S, Saini H, De Fazio S, Carrieri C, Morgan M, Vasiliauskaite L, Benes V, Enright AJ, O'Carroll D. Multiple epigenetic mechanisms and the piRNA pathway enforce LINE1 silencing during adult spermatogenesis. Mol Cell. 2013, 50(4):601–8.

Ellinghaus D, Kurtz S, Willhoeft U. LTRharvest, an efficient and flexible software for de novo detection of LTR retrotransposons. BMC Bioinformatics. 2008, 9:18.

Das PP, Bagijn MP, Goldstein LD, Woolford JR, Lehrbach NJ, Sapetschnig A, Buhecha HR, Gilchrist MJ, Howe KL, Stark R, Matthews N, Berezikov E, Ketting RF, Tavaré S, Miska EA. Piwi and piRNAs act upstream of an endogenous siRNA pathway to suppress Tc3 transposon mobility in the Caenorhabditis elegans germline. Mol Cell. 2008, 31(1):79–90.

Fisher WS. Structure and Functions of Oyster Hemocytes. Immunity in Invertebrates. Springer-Verlag Berlin Heidelberg 1986. ISBN 978-3-642-70770-4. doi:10.1007/978-3-642-70768-1.

Flemr M, Malik R, Franke V, Nejepinska J, Sedlacek R, Vlahovicek K, Svoboda P., A retrotransposon-driven dicer isoform directs endogenous small interfering RNA production in mouse oocytes. Cell. 2013, 155(4):807–16.

Gebert D, Hewel C, Rosenkranz D. unitas: the universal tool for annotation of small RNAs. BMC Genomics. 2017, 18(1):644.

Gebert D, Ketting RF, Zischler H, Rosenkranz D. piRNAs from Pig Testis Provide Evidence for a Conserved Role of the Piwi Pathway in Post-Transcriptional Gene Regulation in Mammals. PLoS One. 2015, 10(5):e0124860.

Grimson A, Srivastava M, Fahey B, Woodcroft BJ, Chiang HR, King N, Degnan BM, Rokhsar DS, Bartel DP. Early origins and evolution of microRNAs and Piwi-interacting RNAs in animals. Nature. 2008, 455(7217):1193–7.

Ha H, Song J, Wang S, Kapusta A, Feschotte C, Chen KC, Xing J. A comprehensive analysis of piRNAs from adult human testis and their relationship with genes and mobile elements. BMC Genomics. 2014, 15:545.

Hirano T, Iwasaki YW, Lin ZY, Imamura M, Seki NM, Sasaki E, Saito K, Okano H, Siomi MC, Siomi H., Small RNA profiling and characterization of piRNA clusters in the adult testes of the common marmoset, a model primate. RNA. 2014, 20(8):1223–37.

Houwing S, Berezikov E, Ketting RF. Zili is required for germ cell differentiation and meiosis in zebrafish. EMBO J. 2008, 27(20):2702–11.

Iwasaki YW, Siomi MC, Siomi H. PIWI-Interacting RNA: Its Biogenesis and Functions. Annu Rev Biochem. 2015, 84:405–33.

Jiang H, Wong WH. SeqMap: mapping massive amount of oligonucleotides to the genome. Bioinformatics. 2008, 24(20):2395–6.

Johnson LS, Eddy SR, Portugaly E., Hidden Markov model speed heuristic and iterative HMM search procedure. BMC Bioinformatics. 2010, 11:431.

Jones BC, Wood JG, Chang C, Tam AD, Franklin MJ, Siegel ER, Helfand SL. A somatic piRNA pathway in the Drosophila fat body ensures metabolic homeostasis and normal lifespan. Nat Commun. 2016, 7:13856.

Kawaoka S, Izumi N, Katsuma S, Tomari Y. 3' end formation of PIWI-interacting RNAs in vitro. Mol Cell. 2011, 43(6):1015–22.

Kiuchi T, Koga H, Kawamoto M, Shoji K, Sakai H, Arai Y, Ishihara G, Kawaoka S, Sugano S, Shimada T, Suzuki Y, Suzuki MG, Katsuma S. A single female-specific piRNA is the primary determiner of sex in the silkworm. Nature. 2014, 509(7502):633–6.

Lagesen K, Hallin P, Rødland EA, Staerfeldt HH, Rognes T, Ussery DW. RNAmmer: consistent and rapid annotation of ribosomal RNA genes. Nucleic Acids Res. 2007, 35(9):3100–8.

Lau YT, Sussman L, Pales Espinosa E, Katalay S, Allam B., Characterization of hemocytes from different body fluids of the eastern oyster Crassostrea virginica. Fish Shellfish Immunol. 2017, 71:372–379.

Lewis SH, Quarles KA, Yang Y, Tanguy M, Frézal L, Smith SA, Sharma PP, Cordaux R, Gilbert C, Giraud I, Collins DH, Zamore PD, Miska EA, Sarkies P, Jiggins FM. Pan-arthropod analysis reveals somatic piRNAs as an ancestral defence against transposable elements. Nat Ecol Evol. 2018, 2(1):174–181.

Li XZ, Roy CK, Dong X, Bolcun-Filas E, Wang J, Han BW, Xu J, Moore MJ, Schimenti JC, Weng Z, Zamore PD. An ancient transcription factor initiates the burst of piRNA production during early meiosis in mouse testes. Mol Cell. 2013, 50(1):67–81.

Lim RS, Anand A, Nishimiya-Fujisawa C, Kobayashi S, Kai T., Analysis of Hydra PIWI proteins and piRNAs uncover early evolutionary origins of the piRNA pathway. Dev Biol. 2014, 386(1):237–51.

Lowe TM, Chan PP. tRNAscan-SE On-line: integrating search and context for analysis of transfer RNA genes. Nucleic Acids Res. 2016, 44(W1):W54–7.

Madison-Villar MJ, Sun C, Lau NC, Settles ML, Mueller RL., Small RNAs, from a Big Genome: The piRNA Pathway and Transposable Elements in the Salamander Species Desmognathus fuscus. J Mol Evol. 2016, 83(3–4):126–136.

Manakov SA, Pezic D, Marinov GK, Pastor WA, Sachidanandam R, Aravin AA. MIWI2 and MILI Have Differential Effects on piRNA Biogenesis and DNA Methylation. Cell Rep. 2015, 12(8):1234–43.

Aravin AA. MIWI2 and MILI Have Differential Effects on piRNA Biogenesis and DNA Methylation. Cell Rep. 2015, 12(8):1234–43.

Miesen P, Girardi E, van Rij RP., Distinct sets of PIWI proteins produce arbovirus and transposon-derived piRNAs in Aedes aegypti mosquito cells. Nucleic Acids Res. 2015, 43(13):6545–56.

Murchison EP, Kheradpour P, Sachidanandam R, Smith C, Hodges E, Xuan Z, Kellis M, Grützner F, Stark A, Hannon GJ., Conservation of small RNA pathways in platypus. Genome Res. 2008, 18(6):995–1004.

Nandi S, Chandramohan D, Fioriti L, Melnick AM, Hébert JM, Mason CE6, Rajasethupathy P, Kandel ER., Roles for small noncoding RNAs in silencing of retrotransposons in the mammalian brain. Proc Natl Acad Sci U S A. 2016, pii: 201609287.

Palakodeti D, Smielewska M, Lu YC, Yeo GW, Graveley BR., The PIWI proteins SMEDWI-2 and SMEDWI-3 are required for stem cell function and piRNA expression in planarians. RNA. 2008, 14(6):1174–86.

Perrat PN, DasGupta S, Wang J, Theurkauf W, Weng Z, Rosbash M, Waddell S., Transposition-driven genomic heterogeneity in the Drosophila brain. Science. 2013, 340(6128):91–5.

Pezic D, Manakov SA, Sachidanandam R, Aravin AA. piRNA pathway targets active LINE1 elements to establish the repressive H3K9me3 mark in germ cells. Genes Dev. 2014, 28(13):1410–28.

Praher D, Zimmermann B, Genikhovich G, Columbus-Shenkar Y, Modepalli V, Aharoni R, Moran Y, Technau U. Characterization of the piRNA pathway during development of the sea anemone Nematostella vectensis. RNA Biol. 2017, 14(12):1727–1741.

Price AL, Jones NC, Pevzner PA. De novo identification of repeat families in large genomes. Bioinformatics. 2005, 21 Suppl 1:i351–8.

Reuter M, Berninger P, Chuma S, Shah H, Hosokawa M, Funaya C, Antony C, Sachidanandam R, Pillai RS. Miwi catalysis is required for piRNA amplification-independent LINE1 transposon silencing. Nature. 2011, 480(7376):264–7.

Roovers EF, Rosenkranz D, Mahdipour M, Han CT, He N, Chuva de Sousa Lopes SM, van der Westerlaken LA, Zischler H, Butter F, Roelen BA, Ketting RF. Piwi proteins and piRNAs in mammalian oocytes and early embryos. Cell Rep. 2015, 10(12):2069–82.

Rosenberg G. A New Critical Estimate of Named Species-Level Diversity of the Recent molluska. American Malacological Bulletin 2014, 32(2):308–322.

Rosenkranz D1, Zischler H. proTRAC - a software for probabilistic piRNA cluster detection, visualization and analysis. BMC Bioinformatics. 2012, 13:5.

Rosenkranz D, Han CT, Roovers EF, Zischler H, Ketting RF. Piwi proteins and piRNAs in mammalian oocytes and early embryos: From sample to sequence. Genom Data. 2015, 5:309–13.

Rosenkranz D, Rudloff S, Bastuck K, Ketting RF, Zischler H., Tupaia small RNAs provide insights into function and evolution of RNAi-based transposon defense in mammals. RNA. 2015, 21(5):911–22.

Russell S, Patel M, Gilchrist G, Stalker L, Gillis D, Rosenkranz D, LaMarre J. Bovine piRNA-like RNAs are associated with both transposable elements and mRNAs. Reproduction. 2017, 153(3):305–318.

Sasaki T, Shiohama A, Minoshima S, Shimizu N., Identification of eight members of the Argonaute family in the human genome. Genomics. 2003, 82(3):323–30.

Schorn AJ, Gutbrod MJ, LeBlanc C, Martienssen R. LTR-Retrotransposon Control by tRNA-Derived Small RNAs. Cell. 2017, 170(1):61–71.e11.

Seto AG, Kingston RE, Lau NC., The coming of age for Piwi proteins. Mol Cell. 2007, 26(5):603–9.

Thomson T, Lin H., The biogenesis and function of PIWI proteins and piRNAs: progress and prospect. Annu Rev Cell Dev Biol. 2009, 25:355–76.

Vourekas A, Zheng Q, Alexiou P, Maragkakis M, Kirino Y, Gregory BD, Mourelatos Z., Mili and Miwi target RNA repertoire reveals piRNA biogenesis and function of Miwi in spermiogenesis. Nat Struct Mol Biol. 2012, 19(8):773–81.

Xu F, Wang X, Feng Y, Huang W, Wang W, Li L, Fang X, Que H, Zhang G. Identification of conserved and novel microRNAs in the Pacific oyster Crassostrea gigas by deep sequencing. PLoS One. 2014, 9(8):e104371.

Zhang Z, Xu J, Koppetsch BS, Wang J, Tipping C, Ma S, Weng Z, Theurkauf WE, Zamore PD. Heterotypic piRNA Ping-Pong requires qin, a protein with both E3 ligase and Tudor domains. Mol Cell. 2011, 44(4):572–84.

Zhang P, Kang JY, Gou LT, Wang J, Xue Y, Skogerboe G, Dai P, Huang DW, Chen R, Fu XD, Liu MF, He S. MIWI and piRNA-mediated cleavage of messenger RNAs in mouse testes. Cell Res. 2015, 25(2):193–207.

Zhao X, Yu H, Kong L, Liu S, Li Q., High throughput sequencing of small RNAs transcriptomes in two Crassostrea oysters identifies microRNAs involved in osmotic stress response. Sci Rep. 2016, 6:22687.

Zhou Z, Wang L, Song L, Liu R, Zhang H, Huang M, Chen H. The identification and characteristics of immune-related microRNAs in haemocytes of oyster Crassostrea gigas. PLoS One. 2014,; 9(2):e88397.

